# Disentangling cadherin-mediated cell-cell interactions in collective cancer cell migration

**DOI:** 10.1101/2021.05.07.442718

**Authors:** Themistoklis Zisis, David B. Brückner, Tom Brandstätter, Joseph d’Alessandro, Angelika M. Vollmar, Chase P. Broedersz, Stefan Zahler

**Affiliations:** Ludwig-Maximilians-University Munich, Department of Pharmacy, Center for Drug Research, 81377 Munich, Germany; Arnold Sommerfeld Center for Theoretical Physics and Center for NanoScience, Department of Physics, Ludwig-Maximilians-University Munich; Institut Jacques Monod (IJM), CNRS UMR 7592 and Université de Paris, 75013, Paris, France; Department of Physics and Astronomy, Vrije Universiteit Amsterdam, 1081 HV Amsterdam, The Netherlands

## Abstract

Cell dispersion from a confined area is fundamental in a number of biological processes, including cancer metastasis. To date, a quantitative understanding of the interplay of single cell motility, cell proliferation, and intercellular contacts remains elusive. In particular, the role of E- and N-Cadherin junctions, central components of intercellular contacts, is still controversial. Combining theoretical modeling with *in vitro* observations, we investigate the collective spreading behavior of colonies of human cancer cells (T24). Inhibition of E- and N-Cadherin junctions decreases colony spreading and average spreading velocities, without affecting the strength of correlations in spreading velocities of neighboring cells. Based on a biophysical simulation model for cell migration, we show that the behavioral changes upon disruption of these junctions can be explained by reduced repulsive excluded volume interactions between cells. This suggests that cadherin-based intercellular contacts sharpen cell boundaries leading to repulsive rather than cohesive interactions between cells, thereby promoting efficient cell spreading during collective migration.

---

Collective cell migration is central to a number of key physiological processes, including morphogenesis during development [1], as well as immune response [2], wound repair [3] and tissue homeostasis [4] in the developed organism. Aberrant cell migration is associated with several pathologies, such as the spread of malignant cancer cells to previously healthy tissues during metastasis [5]. The migratory dynamics of cell collectives in these processes are not merely the outcome of many independently moving cells: they are controlled by cell-cell interactions [6, 7]. Specifically, cells form mechanosensitive cell–cell adhesion junctions (adherens junctions) and coordinate their movements by actively interacting with each other [8]. These interactions facilitate a coordination of collective behavior where a colony of cells invades an empty area [9]. However, it remains unclear how different types of cell-cell interactions control such collective spreading behavior.

The trajectories of single migrating cells are well described by stochastic trajectory models, both for cells migrating on 2D surfaces [10-12] and in confining environments [13-15]. Yet, it is challenging to describe the stochastic collective migration of a cancer cell colony, as cell division and cell-cell contacts have to be taken into consideration. Cell-cell contacts lead to a variety of interactions between cells. Firstly, cells exhibit excluded volume (EV) interactions, where an individual cell occupies space and exerts a repelling force on other cells that move within this space [16]. Secondly, many cell types have the tendency to reorient their direction of motion and move apart upon contact, which is referred to as Contact Inhibition of Locomotion (CIL) [17, 18]. In physical stochastic trajectory models, these interactions are frequently incorporated as a combination of *repulsive interactions*, modelling EV, and *velocity interactions* including velocity alignment as well as CIL [19-23]. Conceptually, there is a key difference between these interactions: while repulsive interactions depend on the relative positions of cells, velocity interactions depend on their motion, i.e. their velocities or polarities. However, it remains unclear how changes in cell-cell contacts within a migrating colony influence these distinct types of interactions and the resulting collective migratory behavior.

Intercellular interactions are strongly dependent on Cadherins, highly conserved calcium-dependent transmembrane proteins that constitute the main component of adherens junctions. Type I classical cadherins (including epithelial (E) and neuronal (N) cadherin as well as P-, R- and M-cadherin [24]) form strong cell–cell adhesion by predominantly homotypic interaction between their extracellular domains [25]. The intracellular cadherin domains connect to β- and α-catenins that associate with the actin cytoskeleton to mediate mechanotransduction [26]. Changes in the normal expression levels of the different cadherin types has been associated with carcinogenesis. One of the most studied processes related to several epithelial tumors is the cadherin switch observed during Epithelial-Mesenchymal transition (EMT). EMT involves the loss of epithelial cell polarity and cell-cell adhesion and the gain of migratory and invasive properties, resulting in the predominance of a mesenchymal phenotype [27]. More specifically, there typically is a strong downregulation of E-Cadherin in parallel with an upregulation of N-Cadherin in EMT. As a result, E-Cadherin adherens junctions disassociate while N-Cadherin junctions establish a relatively weak (compared to E-Cadherin) adherens junction [28].

However, the role of E- or N-Cadherin-mediated intercellular adhesions in cancer cell migration remains controversial. On the one hand, E-Cadherin downregulation has been related to cancer development [29, 30], and it has been shown that the presence of E-Cadherin induces a spreading cell monolayer to retract and form a spheroid aggregate, a process called dewetting [31], suggesting its role as a potent tumor suppressor. On the other hand, a number of studies suggest the opposite effect: E-Cadherin is required for metastasis in multiple models of breast cancer [32], it promotes expansion of bladder carcinoma in situ [33], and is highly present in patients with prostate cancer [34], ovarian cancer [35], and glioblastoma [36]. A similar controversy characterizes the involvement of N-Cadherin in migratory behavior. Although N-cadherin is a marker of EMT and its expression has been associated with the development of multiple cancer types [28], there are studies pointing in the opposite direction. In fact, N-cadherin loss was associated with increased tumor incidence [37] and metastasis [38]. Consequently, a question is yet to be answered: what is the distinct contribution of E- and N-Cadherin junctions to cell-cell interactions and the resulting spreading dynamics of cancer cell colonies?

Here, we aim to investigate this question by combining experimental observations on collectively migrating cells and a minimal physical model of the spreading behavior. We use an epithelial bladder cancerous cell line (T24), which is characterized by high N-Cadherin expression and limited [39] or zero functional levels of E-Cadherin [40, 41]. After initial confinement of a colony of cells to a circular micropattern, the cells are released using chemical tools [42, 43]. We quantify the collective migration by identifying and tracking the entire ensemble of single cell trajectories in each colony. To investigate the effect of cell-cell contacts for the migration, we inhibit E- or N-Cadherin junctions via specific blocking antibodies. In both cases, our analysis reveals that such inhibition leads to a reduced spreading velocity of the cell colonies. To elucidate these dynamics, we develop a minimal active particle model for collective migration, that includes cell proliferation as well as repulsive and CIL interactions. This model shows that inhibiting E- or N-Cadherin has an effect akin to reducing the strength of repulsive cell-cell interactions in the model. In other words, disturbing either of these cadherin junctions decreases the displacement generated when neighboring cells push each other away in order to create space for themselves. Therefore, we show that both E- and N-Cadherins contribute to the maintenance of intercellular contacts that facilitate cell spreading via repulsive interactions, causing cells to move further away from each other. This could be a consequence of cadherins ‘sharpening’ cellular boundaries, through e.g. shape regulation, changes in interfacial tension, or increased cell-cell recognition [44]. These observations indicate the important role of cadherins in metastatic events and their potential as cancer treatment targets.

## Release from a micropatterned circular adhesive area leads to collective cell spreading

To generate an experimental setup for tracking collective cell spreading dynamics, we develop a micropatterned platform from which cells can be released in a standardized manner. Specifically, we design a new patterning approach based on a novel sequence of surface plasma treatment, standard microcontact printing, fibronectin coating and click chemistry steps. This process results in the production of circular fibronectin-coated adhesive areas that are surrounded by cell-repellent azido (PLL-g-PEG) (APP)-coated surfaces. These non-adhesive surfaces can then be activated on demand, via a biocompatible click chemistry reaction between the azide groups of the APP on the surface and added BCN-RGD peptides to allow time-controlled cell migration outside the circular areas [45] (see Materials and Methods and Fig. 1). Subsequently, we use T24 urothelial bladder carcinoma epithelial cells which is a well-established malignant cell line [46], widely used in cell migration research [47-50] and in EMT transition [50]. The cells are detectable using fluorescence microscopy imaging via their nuclear H2B-GFP fluorescent tag.

**Figure 1.**
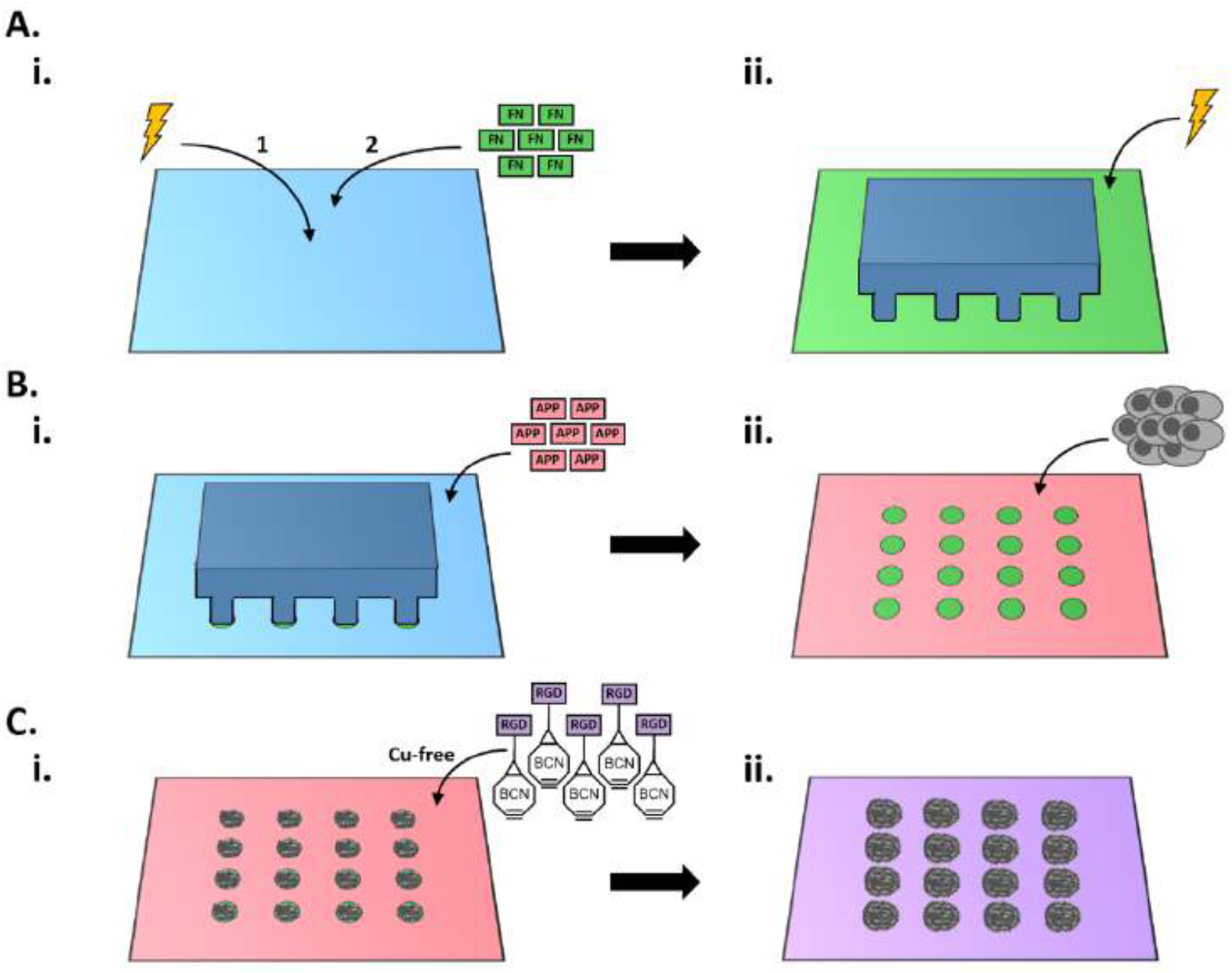
Schematic representation of the microcontact printing and click-chemistry process. A i) Ibidi’s uncoated surface (here one well is represented) undergoes plasma treatment to become reactive, for subsequent attachment of fibronectin (FN). ii) Example of PDMS-square stamp with circular patterns produced with standard microcontact printing techniques (blue). The stamp is placed at the center of the well and the surface is plasma-treated again. The whole surface except for the stamp-protected circular areas loses its fibronectin coating and becomes hydroxylated. B i) With the stamp remaining in place, APP is added next to it and absorbed by the whole surface except for the stamp-protected circular areas (green). B ii) This results in fibronectin-coated circular areas (green) surrounded by an otherwise cell repellent APP surface (red). T24 cancer cells are seeded on the circular areas forming the initial cell population. C i) BCN -RGD peptides are then added and bind to the APP coated surface via click chemistry reaction between the BCN and the azide groups of the APP. C ii) The previously cell-repellent surface is now coated with RGD and thus, highly cell adhesive. The cells are able to expand (migrate) from the circular areas to the rest of the surface.

We perform time-lapse fluorescence and bright-field microscopy for the first 24 hours after surface activation. Here, we observe cells increasingly spreading outwards over time, in all directions, covering a large circular area (Fig. 2 A). While the cells form an approximately confluent monolayer, there are occasional gaps within the layer and significant nearest-neighbor rearrangements during the spreading process (Supplementary Movie S1). Thus, to gain access to the dynamics of the entire cell collective, we perform tracking of the fluorescently tagged nuclei as previously described [43], giving access to the full ensemble of cell trajectories in each escaping cluster (Fig. 2 B).

**Figure 2.**
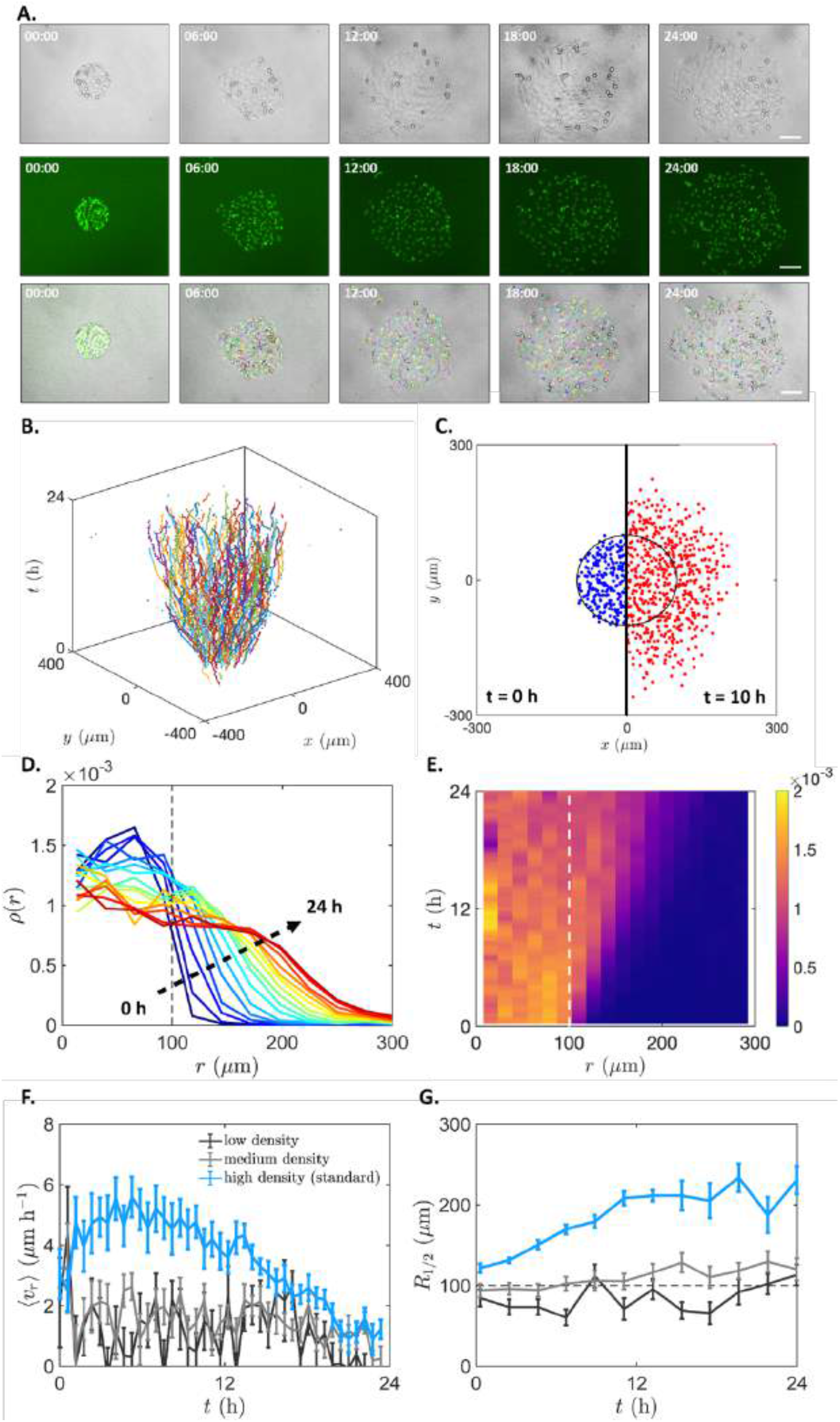
Cell spreading and evolution of cell density of control (untreated) T24 cells. A) Time-lapse bright-field (upper), fluorescence (middle) microscopy images or overlay with cell tracks (lower) showing the T24 cell migration with 6h intervals from 0h to 24h after surface activation. B) Space diagram of colony spreading up to 24h after surface activation. C) Colony spreading radius of T24 cells at 0h (blue) vs 10h (red) after surface activation. D) Evolution of the density profiles over 24 hours (blue to red) plotted as the mean of all colonies (n=14). All curves are separated by 1 h intervals. E) Kymograph of the cell density evolution, corresponding to (D). F) Mean radial velocity (u_r_) over time (average of all colonies per density). The high-density colonies (blue) exhibited a direct increase in radial velocity, larger than the medium and low-density colonies (gray and black, respectively), peaking around 5h after surface activation. G) Average distance where density has decayed to half of its value in the center of the original confinement (i.e. at r=0). The distance was higher over time for the high density compared to the medium and low-density condition. Error bars: SEM; n_high_= 12, n_medium_= 15, n_low_= 12. Space diagram of colony spreading for the different cell densities and complete density evolution profiles shown as mean of all colonies are provided in Supplementary Fig. S4.

At the single cell level, these trajectories are also highly stochastic, as expected from single cells which perform stochastic persistent random motion on unstructured 2D substrates [12]. As shown by the space-time trajectories of the system, the cells have an overall tendency to escape the cluster, and after a period of 10h, a large fraction of the cells has left the initial confinement (Fig. 2 C). The spreading process is quantified by the evolution of the radial density profile *ρ(r)* of the cluster (Fig. 2 D, E). Specifically, we calculate the average number of cells per area element as a function of the distance to the center of the initial confinement radius. As a function of time, the density within the confinement initially decreases, due to cells leaving the confinement through random migration. Correspondingly, the density outside the confinement increases. Interestingly, after a period of approximately 10h, the density inside the confinement stabilizes at a constant value. To further quantify the overall spreading, we calculate the average radius at which the density profile has decayed to half its value at the center of the initial confinement *R*1/2 (Fig. 2 G). Finally, we quantify the average radial velocity of the spreading cells as a function of time, which reveals a marked peak at intermediate spreading times (Fig. 2 F). These statistics are helpful to investigate the impact of collective effects. Thus, we analyzed clusters initialized in the same confinement radius, but with lower cell concentrations. At these lower concentrations, less spreading is achieved (Fig. 2 G), and the peak in radial velocity disappears (Fig. 2 F), indicating that the dynamics observed in our experiments are density dependent, and therefore have a distinct collective character. The collective escape behavior is therefore likely determined by a combination of single-cell motility, cell proliferation, and cell-cell interactions.

## Minimal active particle model captures experimental colony spreading

To elucidate the interplay of the various factors affecting the collective migration in our experiments, we develop a minimal active particle model for collective cell migration (Fig. 3 A). In this model, single cells perform persistent random motion, as observed for single cell migration on two-dimensional substrates [10]. We include cell-cell interactions in our model through two distinct contributions [21, 43, 51]: a repulsive component modelling excluded volume (EV) interactions, and Contact Inhibition of Locomotion (CIL) which models the tendency of cells to reorient away from contacts upon collision. We first confine the particles into a circular region of radius *R* and then observe their behavior upon release, exactly like in the experiment (Methods Section and Fig. 3 B).

**Figure 3.**
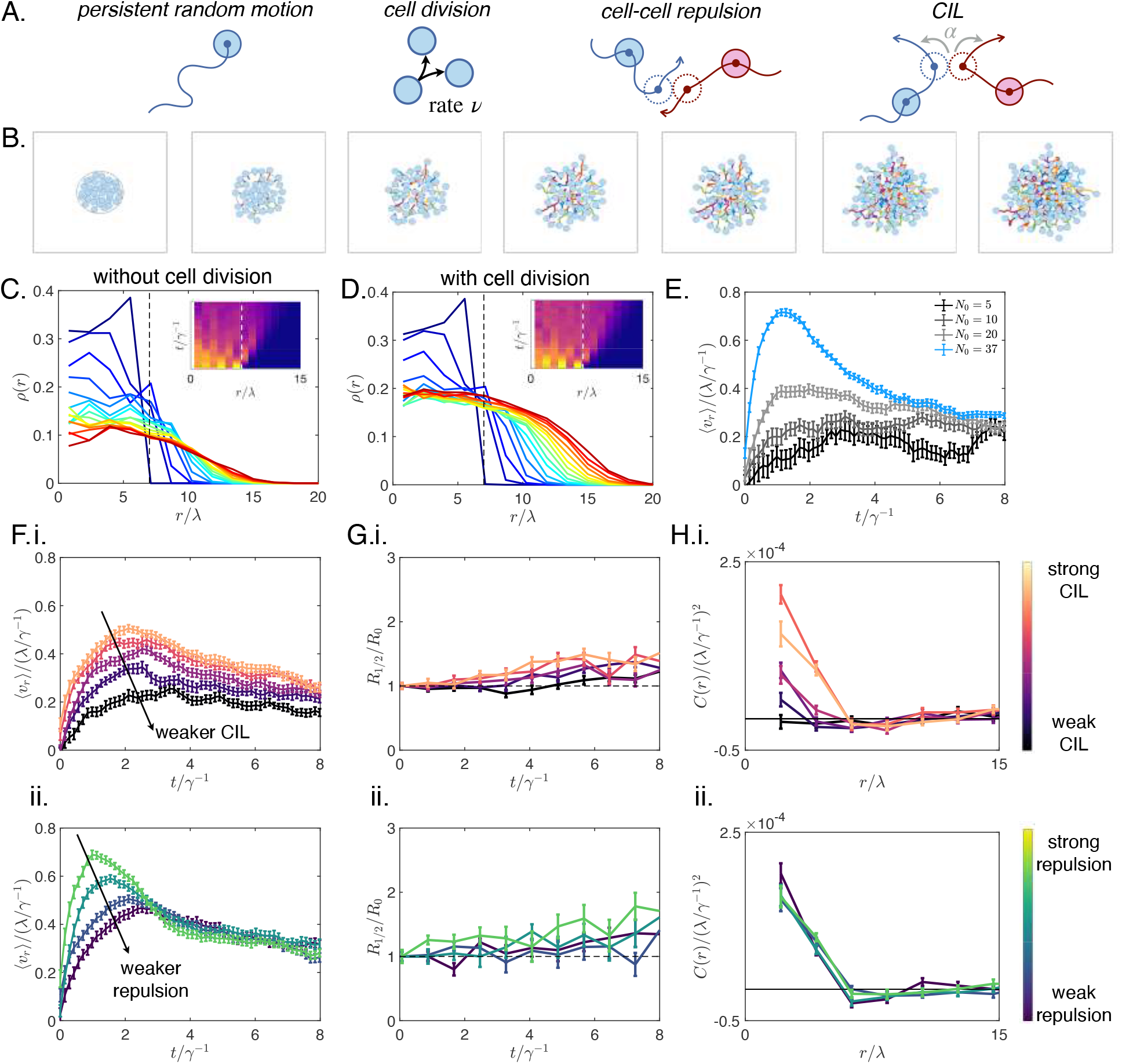
Computational model for collective cell escape. A) Schematic of the components of our active particle model, from left to right: persistent random motion of individual particles, cell division with constant rate *v*, excluded volume interactions, and Contact Inhibition of Locomotion. B) Time-series of a cluster escape simulation. Cell positions are shown as blue circles of radius λ, which is the radius of the repulsive potential. Previous motion of the cells is shown as colored trajectories. C, D) Evolution of the density profile over time (blue to red) plotted as the mean of n=30 colonies. Inset: Kymograph of the cell density evolution. Dashed lines indicate the initial confinement radius. B corresponds to a model without cell division, while C includes cell division. E) Mean radial velocity over time for clusters with different initial density. F) Mean radial velocity over time for clusters with (i) different CIL interaction amplitudes, and (ii) different strengths of cell-cell repulsion interactions. G) Average distance where density has decayed to half of its value in the center of the original confinement (i.e. at r=0). H) Cross-correlation of velocity fluctuations. Error bars: SEM; n=30 for all panels.

Interestingly, this model predicts a rapid decay of the density within the initial confinement area over time, as particles perform random motion and are repelled by their neighbors and move outwards (Fig. 3 C). This observation is inconsistent with our experimental data, which showed only a weak decay in the initial confinement area (Fig. 2 D, E). As shown by our cell proliferation estimations, cell division plays an important role on the time-scale of tissue spreading in this system: the number of cells nearly doubles within 10h (Supplementary Fig. S3). We therefore include a basic implementation of cell division in our model, where cells stochastically perform divisions at a constant rate. This model including cell division exhibits a slower decay of density, and an overall density profile that is consistent with our experimental observations (Fig. 2 D, 3 D). This also suggests that divisions play an important role in the experiment by keeping the cell layer close to confluent. This prevents the density from decreasing too quickly, in which case cells would not interact significantly, further supporting the important role of cell proliferation in collective cell spreading phenomena.

Having included cell division, we find that our model captures other key features of the experimentally observed dynamics. Importantly, we observed that the model predicts a peak in the radial velocity (Fig. 3 F i), similar to experiments (Fig. 2 G). This peak in radial velocity on a time-scale of the order of the persistence time of the cells corresponds to the outward diffusive flux expected for a collection of self-propelled particles [43, 52]. Specifically, upon removal of the confinement, cells at the boundary of the cluster are repelled by the bulk of the cluster, leading to a re-orientation of their movement in an outward direction. This causes the initial increase of the average radial velocity, which is followed by a decreasing trend due to the randomization of movement once the cluster has spread significantly. Furthermore, our model reproduces the gradual increase of the spreading radius (Fig. 3 G i), and a positive cross-correlation of velocity fluctuations indicating short-ranged alignment of cell movement (Fig. 3 H i). Finally, our model correctly predicts a reduction in the radial velocity for lower cell densities, as we observed experimentally (Fig. 3 E). Taken together, these results demonstrate that our cell cluster experiments exhibit the behavior expected for a collection of active particles with interactions. In the experiment, the interactions between cells are known to be controlled by transmembrane proteins, including E- and N-Cadherins [44, 53], whose role we seek to elucidate in the next section.

## Effect of blocking antibody treatment on E- and N-Cadherin gene and protein expression

To investigate the role of E- and N-Cadherin adherens junctions in collective cell migration, we inhibit their function using either E- or N-Cadherin blocking antibodies at different concentrations. To assess the effect of E-Cadherin blocking antibody on the different cadherin gene expression levels, we perform qPCR for E- and N-Cadherin genes at 1h and 5h after E-Cadherin blocking antibody treatment at the highest concentration tested (25 μg/ml). The qPCR serves as a short-term indicator of compensatory reactions of the cells upon functional blocking of an adhesion molecule in the crucial 5h time window after activation. This 5h timepoint coincides with the peak spreading velocities in the control condition and is therefore of particular interest. We find a significant upregulation of the E-cadherin gene expression after 5 hours compared to control (Fig. 4 Ai). This increase can be considered as a compensatory mechanism of the cell to normalize its E-Cadherin functionality after the antibody-mediated blockage. Furthermore, the same treatment results in an early slight upregulation followed by significant downregulation of N-cadherin gene expression at 5h (Fig. 4 Aii). The latter result indicates that the upregulated E-Cadherin blocks the expression of N-cadherin [54, 55], which may correspond to a known phenomenon called cadherin switching (extensively reviewed by Loh et al. [28]). Moreover, using Western Blot (WB), we evaluate the effect of E- or N-blocking antibody on E-Cadherin protein levels, as WB provides a longer time-scale endpoint image of the blocking effect on the total E-Cadherin levels. Here, we observe a significant downregulation of E-Cadherin at 24h after treatment with the E-Cadherin blocking antibody, verifying the antibody functionality. E-Cadherin is also downregulated after N-Cadherin blocking antibody treatment (Fig. 4 B), which further implies the presence of a cadherin switching effect. Specifically, the N-Cadherin blocking antibody could transiently increase the gene expression of N-Cadherin, as a compensatory mechanism, which in turn could represses E-Cadherin expression. Interestingly, for E-Cadherin in the control (untreated) condition, we detect multiple shorter bands rather than one band of 130-135 kDa which is the normal size of the protein. The observed bands were a size of ∼120 kDa, 95 kDa and 55kDa (as shown in Supplementary Fig. S5, respectively). Such deviations from the 135 kDa range, involving predominantly a soluble 80 kDa species [56] [57] [58] as well as 97 kDa [59], 48 kDa [60] and 23 kDa [61] fragments are common in the literature and have been associated with the development of different cancer types [28] [62] [63]. Therefore, as E-Cadherin protein expression is known to be very limited [39] or non-existent [40, 41] in T24 cells, it is probable that the shorter E-Cadherin fragments we see are a result of protein degradation.

**Figure 4.**
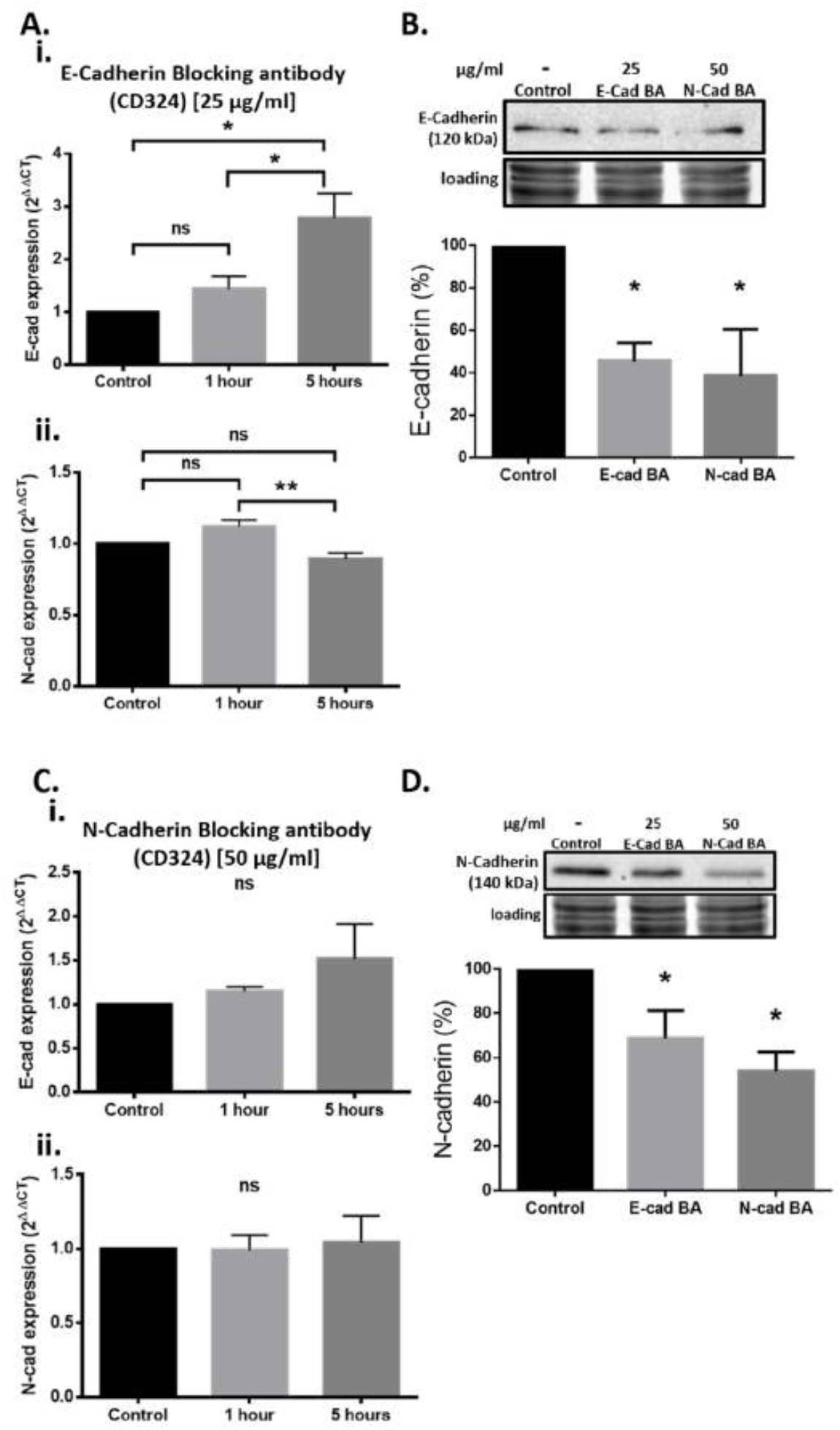
Effect of blocking antibody treatment on E- and N-Cadherin gene and protein expression. A) Quantitative PCR analysis of (i) E- and (ii) N-Cadherin gene expression in untreated (control) or treated T24 cells with E-blocking antibody for 1 and 5 hours, respectively. E-Cadherin blocking antibody treatment at the highest concentration tested (25um/ml) resulted in a significant upregulation of the E-Cadherin gene expression after 5 hours compared to control (mean diff. ± SE= 1.780± 0.4290, p=0.0142). Furthermore, the same treatment resulted in a significant downregulation of N-Cadherin gene expression at the same timepoint compared to the 1h timepoint, indicating a cadherin-switching effect (mean diff ± SE = 0.2251 ±0.05014, p=0.0099). B) Quantitative Western Blot analysis of E-cadherin protein levels in untreated (control) or treated T24 cells with 25μg/ml E-Cadherin or 50μg/ml N-Cadherin blocking antibody after 24 hours. Both antibodies significantly reduced the levels of E-Cadherin after 24 hours (Control vs E-CAD BA: mean diff ± SE=54.43±19.40, p=0.03; Control vs N-CAD BA: mean diff ± SE= 61.47±19.40, p=0.03). C) Quantitative PCR analysis of (i) E- and (ii) N-Cadherin gene expression in untreated (control) or treated T24 cells with N-blocking antibody for 1 and 5 hours, respectively. N-Cadherin blocking antibody treatment at the second highest concentration tested (50 μg/ml) resulted in a non-significant upregulation of N- and E-Cadherin gene expression at 5h compared to control (N-Cadherin: mean diff ± SE= 0.0447±0.1676, p=0.9618; E-Cadherin: mean diff ± SE = 0.5150±0.3258, p=0.3232). D) Quantitative Western Blot analysis of N-Cadherin protein levels in untreated T24 cells (control), cells treated with 25μg/ml E-Cadherin blocking antibody and cells treated with 50μg/ml N-Cadherin blocking antibody. Both antibodies significantly reduced the levels of N-Cadherin after 24 hours (Control vs E-CAD BA: mean diff ± SE=31.37±12.50, p=0.02; Control vs N-CAD BA: mean diff ± SE= 53.80±12.50, p=0.02). Untreated cells were used for data normalization. One representative Western blot is shown per condition including a total protein loading control. Whole Western blots are shown in supplementary Figure S4. Statistical analysis was performed using 1-way ANOVA followed by Tukey’s multiple comparisons (qPCR) or Sidak’s multiple comparisons (WB) test; p< 0.05 (*), p< 0.01 (**); n = 3 (triplicates).

We then investigate the effect of N-Cadherin blocking antibody on cadherin gene expression levels, by performing qPCR for E- and N-Cadherin genes 1h and 5h after N-Cadherin blocking antibody treatment at the second highest concentration tested (50 μg/ml). In that case, a slight tendency towards upregulation of E-Cadherin gene expression is observed at 5h compared to control (Fig 4 Ci), while the N-Cadherin expression levels were not significantly different from untreated cells (Fig. 4 Cii). This lack of significance could result from the fact that in T24 cells, the presence of N-Cadherin is much higher compared to E-Cadherin [39] and thus a higher concentration of blocking antibody would be required for a stronger effect. However, we observe the clear long-term influence of E- or N-blocking antibody on N-Cadherin protein levels by WB where we identify a significant downregulation of N-Cadherin protein levels at 24h after E- and N-Cadherin blocking antibody treatment (Fig. 4 D). Therefore, we conclude that treatment with either E- or N-cadherin blocking antibody starts with a transient upregulation in the corresponding cadherin gene expression which in turn leads to activation of the cadherin switching mechanism that results in the downregulation of the opposite cadherin. This result is further verified by the WB results, where E- or N-cadherin protein levels are significantly downregulated when cells are treated with opposite blocking antibody over the long-term 24h timepoint. With regards to the WB-detected N-Cadherin bands in the untreated condition, a clear band at the expected size (140kDa) is always observed, suggesting that there was no apparent degradation or soluble form as was the case for E-Cadherin. This is not surprising, as N-Cadherin is the predominant and fully functional cadherin in the T24 cell line [39, 64, 65].

In summary, these findings verify that (i) there is a low gene and protein expression of functional (membrane-bound) E-Cadherin in our T24 cells (Supplementary Fig. S5 C, D) and that (ii) besides the direct blocking effect, there is an ‘off target’ blocking effect, where the continuous overexpression of the cadherin being directly blocked leads to a downregulation of the opposite cadherin due to cadherin switching.

## Disrupting E- and N-Cadherin junctions decreases speed of collective spreading

Having quantified the E- and N-Cadherin expression upon different levels of E- or N-Cadherin blocking, we move on to analyzing the collective migration behavior in these conditions. First, we find that a low concentration of E-Cadherin blocking antibody (10 μg/ml) does not significantly affect migration behaviour such as the colony spreading represented by density profiles and radial velocities of the cells (Fig. 5 A, Bi, Ci, Di, Dii and Supplementary Movie S4). However, blocking E-Cadherin at a higher concentration of antibody (25 μg/ml) reduces the average spreading of the colonies (Fig. 5 Bii, Cii, Dii) as well as the average radial velocity of the cells (Fig. 5 Di). Similarly, increasing concentrations of N-cadherin blocking antibody leads to reduced average colony spreading and radial velocities, with the highest one (100 μg/ml) having the strongest effect (Fig. 6 A, B, C, Di, Dii and Supplementary Movie S5). In contrast, we find that the average velocity of single migrating cells in experiments with sparsely seeded cells is not significantly affected by the addition of either blocking antibody, for the whole duration of the experiment (Supplementary Fig S1). Furthermore, the proliferation of cells is similar across all conditions (Supplementary Fig S3). These observations suggest that the change in spreading behaviour upon Cadherin blocking is not mediated by changes in the behaviour of single cells or their proliferation, but is mainly caused by the reduction in cell-cell interactions and is thereby a collective effect.

**Figure 5.**
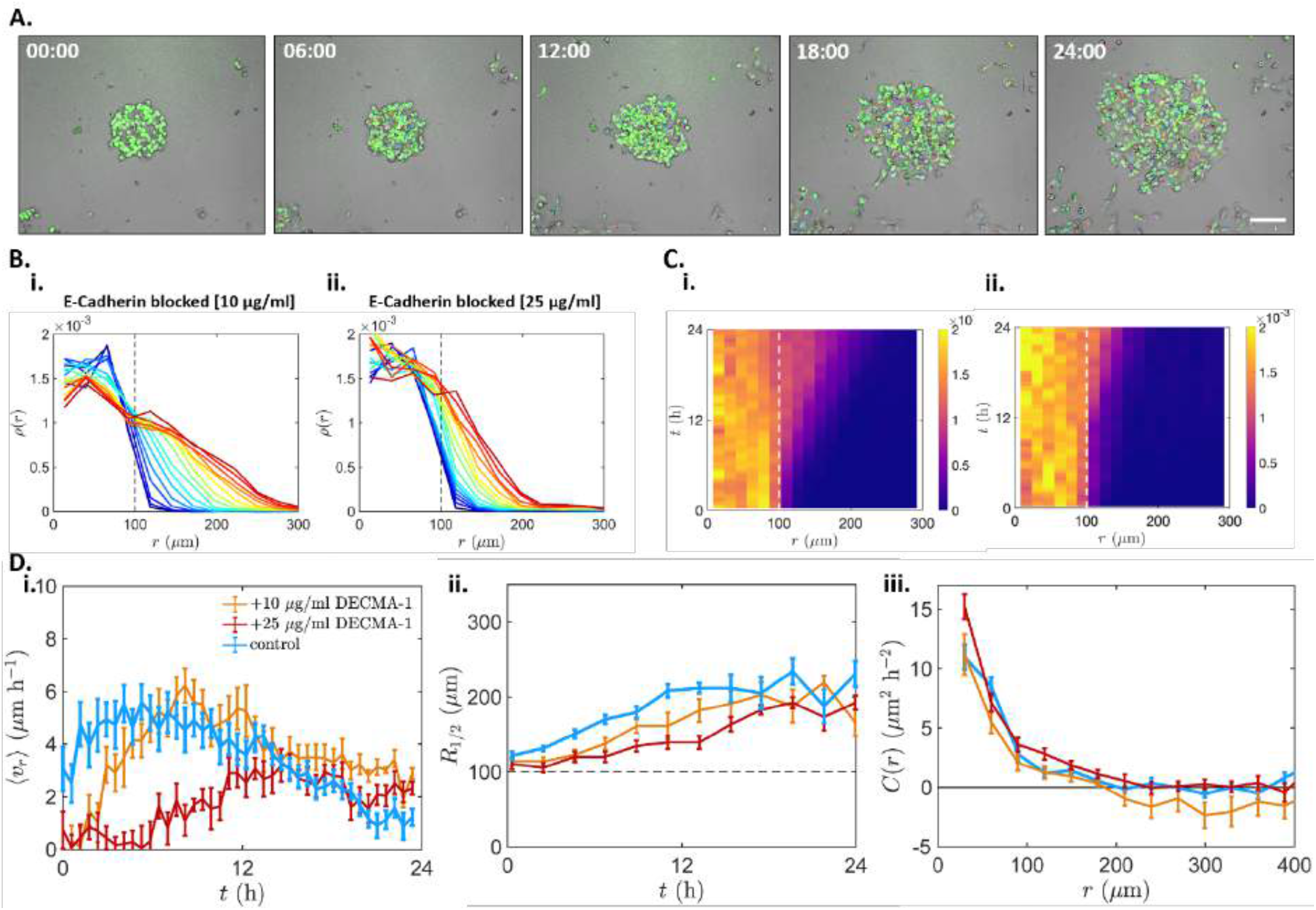
Evolution of cell density profile, radial velocities and average distance of T24 cells treated with increasing concentrations of E-Cadherin blocking antibody. A) Time-lapse overlay of bright-field and fluorescence microscopy images with cell tracks of the 25 μg/ml E-Cadherin blocking, showing the T24 cell migration with 6h intervals from 0h to 24h after surface activation. B) Space diagram of colony spreading up to 24h after surface activation. B) Evolution of the density profiles over 24 hours (blue to red) plotted as the mean of all colonies per condition for T24 cells treated with (i) 10 μg/ml or (ii) 25 μg/ml E-Cadherin blocking antibody. All curves are separated by 1 h intervals. C) Kymographs of the cell density evolution, for T24 cells treated with (i) 10μg/ml and (ii) 25 μg/ml E-Cadherin blocking antibody. D) i) Mean radial velocity (u_r_) over time (average of all colonies per condition). The control condition exhibited a direct increase in radial velocity, peaking around 5h after surface activation (blue). 10 μg/ml E-Cadherin blocking antibody slowed down this increase in radial speed, which peaked at 8h (orange). The highest concentration of blocking antibody (25 μg/ml) resulted in even lower radial velocity that did not reach the initial peaks exhibited in the other conditions (red). ii) Average distance where density has decayed to half of its value in the center of the original confinement (i.e. at r=0). The distance was the highest over time in the control condition and decreased with increasing concentrations of E-Cadherin blocking antibody. iii) Cross correlation of velocity fluctuations showing no significant differences between conditions. Error bars: SEM; n_control_= 12, n_10ECAD_= 13, n_25ECAD_= 8.

**Figure 6.**
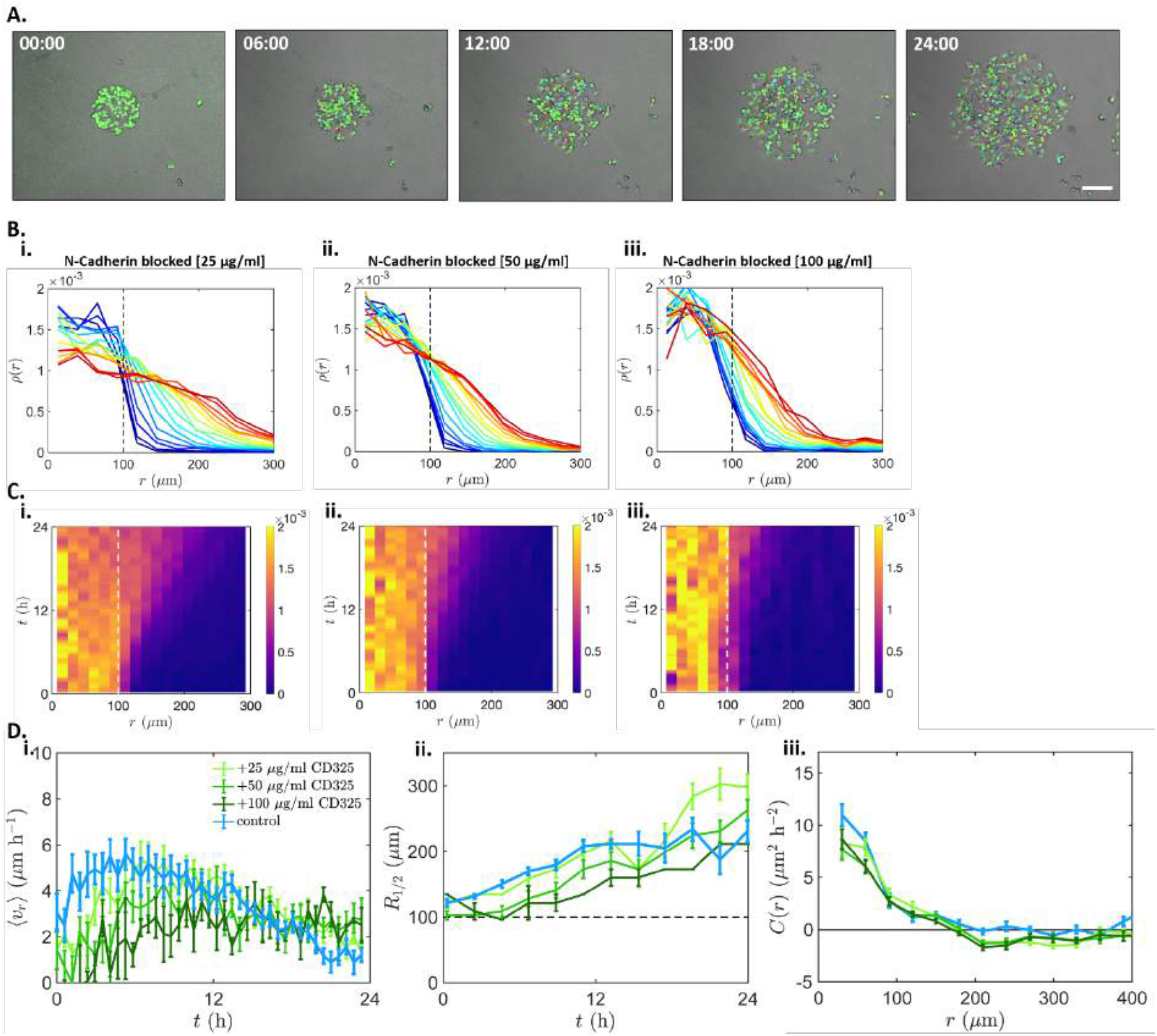
Evolution of cell density profile, radial velocities and average distance of T24 cells treated with increasing concentrations of N-Cadherin blocking antibody. A) Time-lapse overlay of bright-field and fluorescence microscopy images with cell tracks of the 100 μg/ml N-Cadherin blocking, showing the T24 cell migration with 6h intervals from 0h to 24h after surface activation. B) Evolution of the density profiles over 24 hours (blue to red) plotted as the mean of all colonies per condition for T24 cells treated with (i) 25 μg/ml, (ii) 50 μg/ml or (iii) 100 μg/ml N-Cadherin blocking antibody. All curves are separated by 1 h intervals. C) Kymographs of the cell density evolution, for T24 cells treated with (i) 25 μg/ml, (ii) 50 μg/ml or (iii) 100 μg/ml N-Cadherin blocking antibody. D) i) Mean radial velocity (u_r_) over time (average of all colonies per condition). The control condition exhibited a direct increase in radial velocity, peaking around 5h after surface activation (blue). Increasing concentrations of N-Cadherin blocking antibody reduced this increase in radial speed, with the highest reduction observed in the 100μg/ml treated cells (dark green). ii) Average distance where density has decayed to half of its value in the center of the original confinement (i.e. at r=0). The distance was the highest in the control condition and decreased with increasing concentrations of N-Cadherin blocking antibody up to 11h. After this timepoint, the 25 μg/ml N-Cadherin blocking antibody treated colonies surpassed the control ones. iii) Cross correlation of velocity fluctuations showing no significant differences between conditions. Error bars: SEM; n_control_= 12, n_25NCAD_= 8, n_50NCAD_= 6 n_100NCAD_= 3.

To identify a possible change in cell-cell interactions due to cadherin blocking, we calculate the cross-correlation functions of velocity fluctuations between pairs of cells, which quantifies how similar cellular velocities are as a function of their distance from one another (Methods Section and Fig. 5 Diii and 6 Diii). As expected, in the control condition, we find that cells tend to align their direction of motion with neighbouring cells, but exhibit no correlations at long distances. Unexpectedly, however, we find that all observed experimental conditions have a similar cross-correlation function. This indicates that while we expect a change in cell-cell interactions to be responsible for the change in spreading behavior, this change does not directly affect the degree of velocity alignment, quantified through the velocity cross-correlation. In a following section, we will turn to a theoretical model for a possible explanation of these observations.

To summarize our experimental findings, we find that by partially blocking either E- or N-cadherin adherens junctions, the collective spreading behaviour of initially confined clusters of T24 cells becomes less efficient. This suggests that cell-cell contacts are important for coordinated migration, possibly by promoting cell-cell interactions. This result is in agreement with earlier reports showing that preventing cells from forming stable cell-cell contacts resulted in uncoordinated and random cell movement [66], leading to significantly lower migration velocities [67]. In contrast to other studies observing no E-Cadherin expression in T24 cells, we detect its presence (120 kDa) among other fragmented species of the protein. Furthermore, we show that as a type III carcinogenic line, T24 cells exhibit an increased N-Cadherin vs E-Cadherin expression ratio (3/1 as shown in Fig. S 5 C, D), characteristic for EMT [28]. Interestingly, we find that the limited E-Cadherin expression is still important for the efficiency of the collective migration, as is the more predominantly expressed N-Cadherin. Therefore, the interplay between E- and N-Cadherin in T24 cells points to a crucial balance in cell-cell contacts that seems to be important for collective migration. In the next section, we use our minimal active particle model to elucidate the nature of these interactions and how they influence the cell spreading behavior.

## Varying cell-cell interactions in a minimal active particle model captures the effects of Cadherin blocking

To investigate how changes in cell-cell interactions affect the spreading behavior in our model, we first vary the strength of contact inhibition of locomotion (CIL). We implement CIL as an angular repulsion that acts as a torque on cells undergoing a contact, with strength *α*, similar to previous work [43] (see Fig. 3 A and Methods Section). We find that decreasing *α*, corresponding to weaker CIL, leads to a reduction in radial velocity, spreading, and cross-correlations (Fig. 3 F i, G i, H i). Thus, while the first two findings are in line with the changes in behavior upon cadherin inhibition in the experiment, the change in cross-correlation is not observed in the experiment. In contrast, reducing the strength of the repulsive interactions between particles leads to a reduction of the radial velocity peak and the overall spreading, while keeping the cross-correlations constant (Fig. 3 F ii, G ii, H ii) - similar to what we observed experimentally upon blocking E- or N-Cadherin-mediated intercellular contacts (Fig. 5 B ii, D and 6 B ii-iii, D). These results are robust over a wide range of parameters in the model (Supplementary Figs. 8-11). These observations suggest that disrupting cell-cell junctions through E and N-Cadherin blocking has an effect akin to reducing excluded volume interactions between cells.

The reduced spreading for weaker CIL and weaker repulsive interactions can be understood intuitively. Firstly, CIL interactions ensure that cells at the cluster boundary do not cross paths, leading to outward alignment of their velocities. In fact, in this setup, CIL has an effect very similar to velocity alignment interactions: an alternative model with velocity alignment instead of CIL produces very similar results (Supplementary Fig. 7), highlighting the similarity of these two interaction types in this setup. Secondly, repulsion ensures that boundary cells are repelled by the bulk of the cluster, which further rectifies their motion into a radially outward direction. Thus, both stronger CIL and stronger repulsive interactions lead to faster, more efficient spreading dynamics (Fig. 3 F, G).

However, we can distinguish the two types of interaction through the cross-correlation of cell velocities: this quantity serves as a good indicator for changes in CIL-behavior. Specifically, changing repulsive interactions has no significant effect on the correlation function, since it is a position-dependent interaction (Fig. 3 H ii). In contrast, CIL is a velocity-dependent interaction, and its strength therefore controls the magnitude of the velocity cross-correlations (Fig. 3 H i).

Taken together, these results show that cell-cell interactions are key drivers of tissue spreading in this setup, and that disrupting cell-cell junctions through E- and N-Cadherin blocking has an effect akin to reducing repulsive interactions between cells. Therefore, the congruity between our experimental and modeling data suggests that both E- and N-Cadherin-mediated intercellular contacts create repulsive events via excluded volume interactions that are critical for the efficient cell spreading during collective migration. This effect could be due to cadherins ‘sharpening’ cell boundaries by for example regulating cell shape, improving cell-cell recognition, or increasing interfacial tension. Indeed, both E- and N-cadherin have been shown to determine inter-cellular interfacial tension in the developing epithelium [44, 68, 69]. These results are also in qualitative agreement with previous work where the interactions of colliding pairs of cells were inferred directly from observed trajectories [51]. Specifically, it was shown that the cancerous MDA-MB-231 cell line exhibits less repulsive interactions than the non-malignant MCF10A cell line, which is known exhibit higher E-cadherin expression than MDA-MB-231 cells [70, 71]. Our work therefore further supports the important role of cadherin-mediated cell-cell interactions, and elucidates their role in collective cell migration.

This study provides new insight into the role of different cadherin junctions in the dynamics of collective cancer cell migration. In our setup, we reveal that blocking E- or N-Cadherin in collectively migrating T24 cancer cells significantly reduces their spreading efficiency. The observed phenomenology is well captured by a biophysical model of stochastically migrating cells. Our model shows that cell proliferation as well as the excluded volume and Contact Inhibition of Locomotion interactions between cells drive tissue spreading in our setup. Our combined experimental and theoretical results further indicate that disrupting E- and N-Cadherin-mediated intercellular contacts leads to a decrease in repulsive cell-cell interactions, which in turn reduces the spreading efficiency of the cell collective. Therefore, from a biomedical point of view, this study underscores the importance of E- and N-Cadherins as potential pharmacological targets in metastatic cancer research. Furthermore, our experimental setup design could be adapted for future research in the field, such as studying the impact of mechanical cell-cell communication on cell spreading on mechanically compliant substrates [72-74], or chemotactic cell spreading in external gradients [75, 76].

## Methods

### T24 cell culture transfection with H2B-GFP plasmid for nucleus labeling

H2B–GFP expression vectors, were obtained from Addgene (#11680). T24 cells exponentially growing in Dulbecco’s modified Eagle’s medium (DMEM) supplemented with 10% fetal calf serum (FCS) were transfected with 2.5 µg of the H2B–GFP expression vector carrying a G418 resistance as selection marker, using an Amaxa R-Kit (Program I-013) under constant humidity at 37°C and 5% CO2. 24h after the transfection, cells were treated with G418 (A1720, Sigma-Aldrich) to an end concentration of 0.8mg/ml in 2ml well-plates and then further cultivated in T25 flasks and later on in T75 flasks with the same concentration of G418 (0.8mg/ml). After two rounds of additional cell sorting by flow cytometry the GFP+ cells at passage 30 were frozen in a nitrogen tank at a concentration of 1× 10^6^ cells/ml.

For all collective migration experiments, T24 cells were pre-grown as monolayers and diluted down to the desired concentrations in Dulbecco’s modified Eagle’s medium (DMEM) supplemented with 10% fetal calf serum (FCS), 10.000 U/ml penicillin/streptomycin and 0.8 mg/ml antibiotic G418 under constant humidity at 37°C and 5% CO_2_.

### Microcontact printing for circular pattern generation

8-well uncoated μ-Slides (ibidi, Martinsried, Germany) underwent 3 min of oxygen plasma treatment (Plasma cleaner typ ‘‘ZEPTO,’’ Diener electronic, Ebhausen, Germany) at 0.3 mbar for activation (generation of OH-hydroxyl bonds). Then, 250 μl/well of 0.05 mg/ml fibronectin (R&D Systems, US) solution in MilliQ were added to the now highly reactive surface and incubated at room temp for 2 hours. After washing 2 times with 500 μl of milliQ H_2_O the surface was allowed to dry. Following that, we used standard microcontact printing techniques to create PDMS stamps with circular patterns. We placed one stamp at the center of each well and plasma treated the surface one last time at the same conditions as before. This step removes all fibronectin from the surface except the areas that are protected by the stamp, so all the unprotected areas on the surface become hydroxylated and highly reactive again. Without removing the stamps, we added a 7 μl drop of 1mg/ml PLL(20)-g[3.5]-PEG-N_3_(3) (APP) (Susos AG, Switzerland) solution in MilliQ right next to each stamp allowing surface tension to absorb the liquid underneath the stamp. We let the above condition settle for 45 min. We gently removed the stamp and washed 2 times with 500 μl of MilliQ. Now the circular areas contain fibronectin and are highly cell-adhesive while the surrounding areas are initially cell repellent. At this point, T24 cells were trypsinized after reaching confluency, diluted to the desired density (70.000 cell/ml) in the aforementioned DMEM-based medium and 250 μl of this cell suspension were added in each well and allowed to settle overnight at 37 °C. The next day, the cell medium was replaced with 200 μl of fresh medium and the slide was placed under the microscope. Finally, 10 μl of 100μM BCN-cRGDfk (Synaffix, Netherlands) in PBS were added in the medium of each well to a final concentration of 20 μM. The BCN groups formed a link with the Azide groups of the APP-covered, cell-repellent areas around the colonies. This resulted in the binding of RGD on the surface, thereby rendering the surrounding areas cell adhesive and initiating cell migration.

### Blocking antibody treatment of T24 circular colonies

For the blocking antibody treatment experiments, we followed the exact same cell preparation protocol as above with the addition of the following steps: On the next day, after the first washing step, 200 μl of 5mM EGTA solution were added in each well for 30 min. This step was performed in order to break the existing cadherin junctions and allow the blocking antibodies (anti N-Cadherin antibody: LEAF^™^ Purified anti-human CD325, #350804, Biolegend, USA; anti E-Cadherin antibody: CD324 #16-3249-82, Invitrogen, USA) to bind to their respective epitopes. Following that, the wells were washed two times with 200 μl of fresh cell medium. Subsequently, 200 μl of the appropriate E-(10 or 25μg/ml) or N-(25, 50 or 100μg/ml). Cadherin blocking antibody solution in cell medium were added in each well. Cells were incubated additionally for 30 min and then the slide was placed under the microscope. Finally, 10 μl of 100μM BCN-cRGDfk (Synaffix, Netherlands) in PBS were added in the medium of each well to a final concentration of 20 μM.

### Cell imaging

Live cell imaging was performed using the T24 seeded 8-well fibronectin/APP patterned slides with an Eclipse Ti inverted microscope (Nikon, Dusseldorf, Germany) with a 4x/10x phase contrast objective and a CCD camera ([DS-Qi1Mc] Nikon, Dusseldorf, Germany). The slides were inserted into a 37 °C heating and incubation system that was flushed with actively mixed 5% CO_2_ at a rate of 10 l/h, and the humidity was kept at 80% to prevent dehydration. The cells were imaged in bright-field and the fluorescence of the nuclei was detected at a 488 nm wavelength using the integrated fluorescence LED. Time-lapse video microscopy was performed with a time interval of 5 min between images over 24 h.

### Tracking of single cell trajectories

The positions of individual cells were detected as previously described [43] using custom-made ImageJ macros implementing the ‘Find Maxima’ built-in function. The individual trajectories were then reconstructed using a squared-displacement minimization algorithm (http://site.physics.georgetown.edu/matlab) and data analysis was performed via custom-made Matlab programs.

### qPCR

T24 cells were lysed for mRNA isolation. Briefly, “Buffer RLT, Lysis Buffer” (RNeasy^®^ Mini Kit (250) PCR lab) was mixed with DTT 2M at a ratio of 50:1. After medium aspiration and ice-cold PBS rinsing, ice-cold lysis buffer was added and the lysates were stored at -80 °C. For the mRNA, isolation the RNeasy^®^ Mini Kit (250) (QIAGEN, Hilden, Germany) was used according to the modified manufacturer’s instructions. 2 µl of the mRNA samples was used directly for mRNA concentration determination using a Nanodrop^®^ Spectrophotometer (PEQLAB Biotechnologie, Erlangen, Germany) with absorption at 260 nm (specific for mRNA) while impurities were determined at 280 nm. For the reverse transcription of mRNA to cDNA, 2X RT master mix was prepared containing: 10% TaqMan RT Puffer-10x, 0,04% dNTPs, 10% random hexamers, 5% Reverse Transcriptase, 21%RNAase free water, 50% H2O+ RNA 2.5µg. For the quantitative PCR the following primers were obtained from metabion GmbH: E-Cadh_1_F (MM125, 5’
sTGG GCC AGG AAA TCA CAT CC3’), E-Cadh_1_R (MM126, 5’GGC ACC AGT GTC CGG ATT AA3’); N-Cadh_2_F (MM133, 5’CCT TTC AAA CAC AGC CAC GG3’), N-Cadh_2_R (MM134, 5’TGT TTG GGT CGG TCT GGA TG3’). We used 2 µl of the acquired cDNA in each well of the MicroAmp^®^ Fast Optical 96-Well Reaction Plate or 2 µl of autoclaved Millipore H2O for the no-template controls (NTCs), respectively. 10.5 µl of PCR master mix containing 6.25 µl of PowerUPTM SYBR^®^ Green Master Mix, 3.75 µl of autoclaved Millipore H2O, 0.25 µl of forward primer and 0.25 µl of reverse primer were added to each probe well and the qPCR was performed in a QuantStudio^™^ 3 Real-Time PCR system (ThermoFisher). Data were normalized to the housekeeping gene GAPDH. The analysis was carried out with the ΔΔCT method as previously described [77], using the ThermoFisher cloud and threshold cycle was set to > 9-15 and ≤ 30 to allow acceptable detection for best reproducibility.

### Western Blots

Cells were harvested and lysed in RIPA lysis buffer containing a protease inhibitor mix (Roche #4693159001). Lysates were centrifuged at 10,000 x g for 10 min and 4 °C. Protein amounts were assessed by Bradford assay, and an equal amount of protein was separated by SDS-PAGE and transferred to nitrocellulose membranes (Hybond-ECLTM, Amersham Bioscience). Membranes were incubated with blocking buffer containing 5% BSA and 0.1% Tween 20 in PBS for 1h at room temperature, followed by 3x 5 min. rinsing with PBS-T. After that, membranes were incubated with rabbit anti-ECAD (24E10) monoclonal Ab (1:500; #3195, Cell Signaling Technology, Inc. USA) or rabbit anti-NCAD (D4R1H) XP^®^ monoclonal Ab (1:500; #13116, Cell Signaling Technology Inc. USA) at 4°C overnight. Membranes were washed again with PBS-T 3 times for 5 min. Secondary antibody (HRP-Goat-Anti-Rabbit 1:1000; #111-035-144, Dianova, Germany) were used for 2h incubation at room temperature and subsequently conjugated with horseradish peroxidase and freshly prepared ECL solution (protected from light), which contained 2.5 mM luminol (detailed description of ECL solution preparation in table 1). Conjugated proteins were detected by the ChemiDoc^™^ Touch Imaging System (Bio-Rad, USA) and quantified by ImageLab software (Bio-Rad, USA). For quantification protein amount was normalized to total protein-loading, detected by 2,2,2-trichloroethanol activation as described previously [77] [78].

**Table 1.**
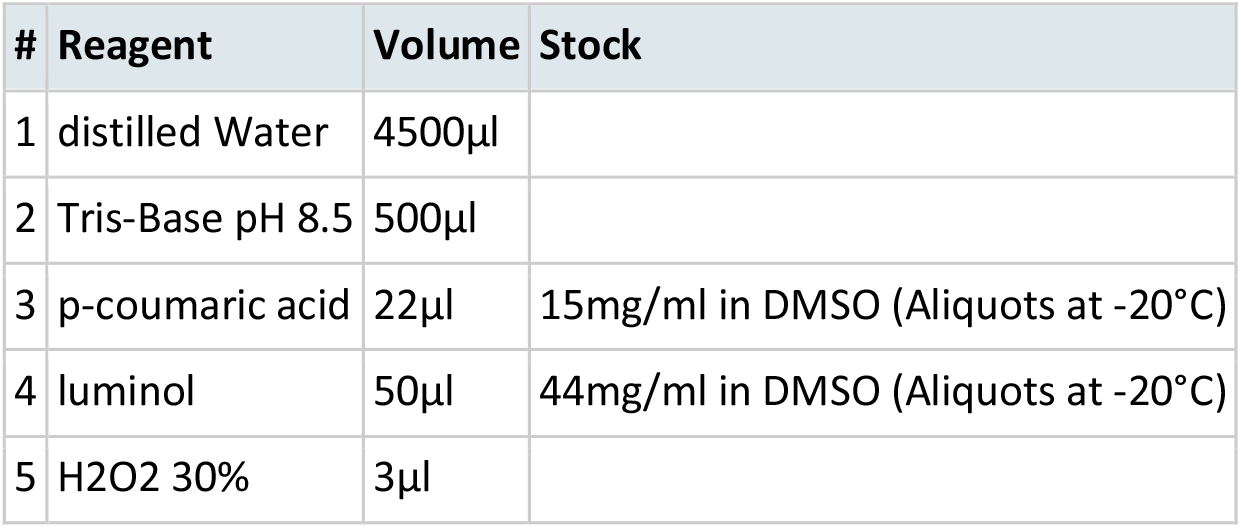
Western Blot Solution Reagents.

| # | Reagent | Volume | Stock |
| --- | --- | --- | --- |
| 1 | distilled Water | 4500µl |  |
| 2 | Tris-Base pH 8.5 | 500µl |  |
| 3 | p-coumaric acid | 22µl | 15mg/ml in DMSO (Aliquots at -20°C) |
| 4 | luminol | 50µl | 44mg/ml in DMSO (Aliquots at -20°C) |
| 5 | H<sub>2</sub>O<sub>2</sub> 30% | 3µl |  |

### Cross-correlation functions of velocity fluctuations

To investigate the interactions of cells in the experiment, we calculate the spatial velocity cross-correlation function

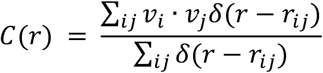

where *v*_*i*_ is the two-dimensional velocity vector of cell *i* and δ(*r* − *r*_*ij*_) is the Dirac delta-function. This function measures how ‘similar’ the velocities (magnitude and direction) of cells at distance *r* from one another are on average. Using discrete bins as an approximation for the delta-function for finite data, we obtain expected results both for experimental and simulated data.

The complete velocity field is composed of the collective outward motion, a dilatational mode, and additional velocity fluctuations due to interactions between the cells. Following previous work [79], we calculate these fluctuations by obtaining the scalar dilatation Λ as a function of time, by optimizing the quantity ∑_*i*_[*x*_*i*_(*t + T*) − Λ*x*_*i*_(*t*)]^7^. The fluctuation velocities are then giving by *u*_*i*_ = [*x*_*i*_(*t i T*) − Λ*x*_*i*_(*t*)]/*T*. Note that here, we use a time-interval *T* = 15Δ*t* which is larger than the time-resolution of the experiment. This allows us to average out the short-time scale noise fluctuations of the cellular velocities, and instead focusses on longer time-scale process relevant to the spreading dynamics. We test this approach in our simulations, and find that it accurately detects the presence of velocity-dependent interactions, such as CIL (Supplementary Figure 6).

### Computational modeling

To provide a minimal computational model for the escape process, we implement a simple active particle model for collective cell migration. Similar to previous works [21, 22, 43, 80-82], we describe the motion of the cells using stochastic equations of motion with interactions. Specifically, we use the equation of motion

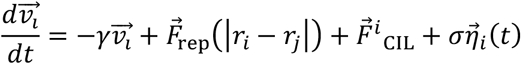

where 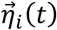 represents a Gaussian white noise with 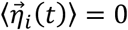 and 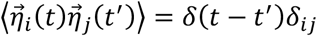. The model furthermore includes a persistence term −*γv*, where *γ*^−1^ is the persistence time of the cells. The repulsive interactions are implemented as the repulsive part of a quadratic potential

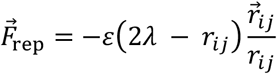

where *λ* represents the radius of the cells, and *ε* is the strength of the interaction.

The contact inhibition of locomotion (CIL) interaction 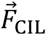 is implemented in the form of a rotation of the velocity vector away from the distance vector 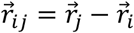 to nearest neighbours, which are defined by being within an interaction range of radius 2.5*λ*, and being on collision course with cell *i*, i.e. 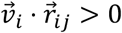. The angular displacement only depends on the velocity direction, a constant acceleration *α* and the number as well as the positions of nearest neighbours: For each nearest neighbour the direction of the axis of rotation is found such that the rotation will be away from the nearest neighbours. All directions of the axes of rotations of all nearest neighbours are added up and multiplied by the acceleration *α*. Specifically, we use

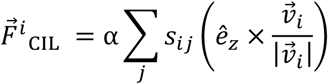

Where

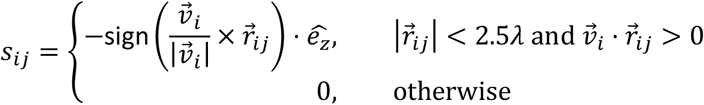

In simulations where velocity alignment rather than CIL is used (Supplementary Figure 8), we replace 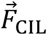 by an alignment interaction 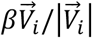 with strength *β*, which is implemented as a constant acceleration in the direction of the average velocity 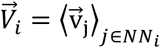of nearest neighbours within an interaction range of radius 2.5*λ*.

Finally, cell division is implemented with a constant probability *vdt* of dividing, provided there is sufficient space for the appearance of new cells. In a division event, a cell produces a daughter cell in its direct neighborhood with an initial velocity pointing away from the mother cell.

The simulation is performed in non-dimensional units such that γ^−1^ = *λ* = 1. We use the parameters σ^7^ = 2, *v* = 0.1, and vary *ε* between 0.1 and 40, and *α* between 0 and 12. We initialize *N* = 37 particles within the initial confinement radius *R*. The stochastic trajectories of the model are then simulated by step-wise Euler updates with a time-step of *dt* = 10^−1^. We first perform a pre-equilibration run with a confinement potential at *r* = *R*, modelling the initial confinement phase. At *t* = 0, we remove the boundary by setting the confinement potential to zero, leading to the escape of the simulated cluster.

### Statistical evaluation

For statistical analysis of the data one-way ANOVA followed by Dunnett’s multiple comparisons test was performed using GraphPad Prism version 8.0.0 for Windows, (GraphPad Software, San Diego, California USA, www.graphpad.com). n.s.= not significant, * p < 0.05, ** p < 0.01.

## Author Contributions

T.Z., D.B.B., C.P.B. and S.Z. designed the study. T.Z. performed all experiments. J.A. contributed tracking software. D.B.B. and T.Z. analyzed data. D.B.B. and T.B. developed the theoretical model. T.Z. and D.B.B. wrote the paper with input from all authors.

## Acknowledgements

Funded by the Deutsche Forschungsgemeinschaft (DFG, German Research Foundation) - Project-ID 201269156 - SFB 1032 (Projects B8 and B12). D.B.B. is supported in part by a DFG fellowship within the Graduate School of Quantitative Biosciences Munich (QBM) and by the Joachim Herz Stiftung.

## Supplementary Information for

E- and N-Cadherin-mediated intercellular contacts enhance collective spreading of migrating cancer cells

### Movie Descriptions

**Supplementary Movie S1:** Video of bright-field microscopy imaging showing the control (untreated) T24 cell migration from 0h to 24h after surface activation.

**Supplementary Movie S2:** Video of fluorescence microscopy imaging showing the control (untreated) T24 cell migration from 0h to 24h after surface activation.

**Supplementary Movie S3:** Overlay video of bright-field and fluorescence microscopy imaging with cell tracks showing the control (untreated) T24 cell migration from 0h to 24h after surface activation.

**Supplementary Movie S4:** Overlay video of bright-field and fluorescence microscopy imaging with cell tracks showing the T24 cell migration upon 25 μg/ml E-Cadherin blocking, from 0h to 24h after surface activation.

**Supplementary Movie S5:** Overlay video of bright-field and fluorescence microscopy imaging with cell tracks showing the T24 cell migration upon 100 μg/ml N-Cadherin blocking, from 0h to 24h after surface activation.

---

**Figure S1.**
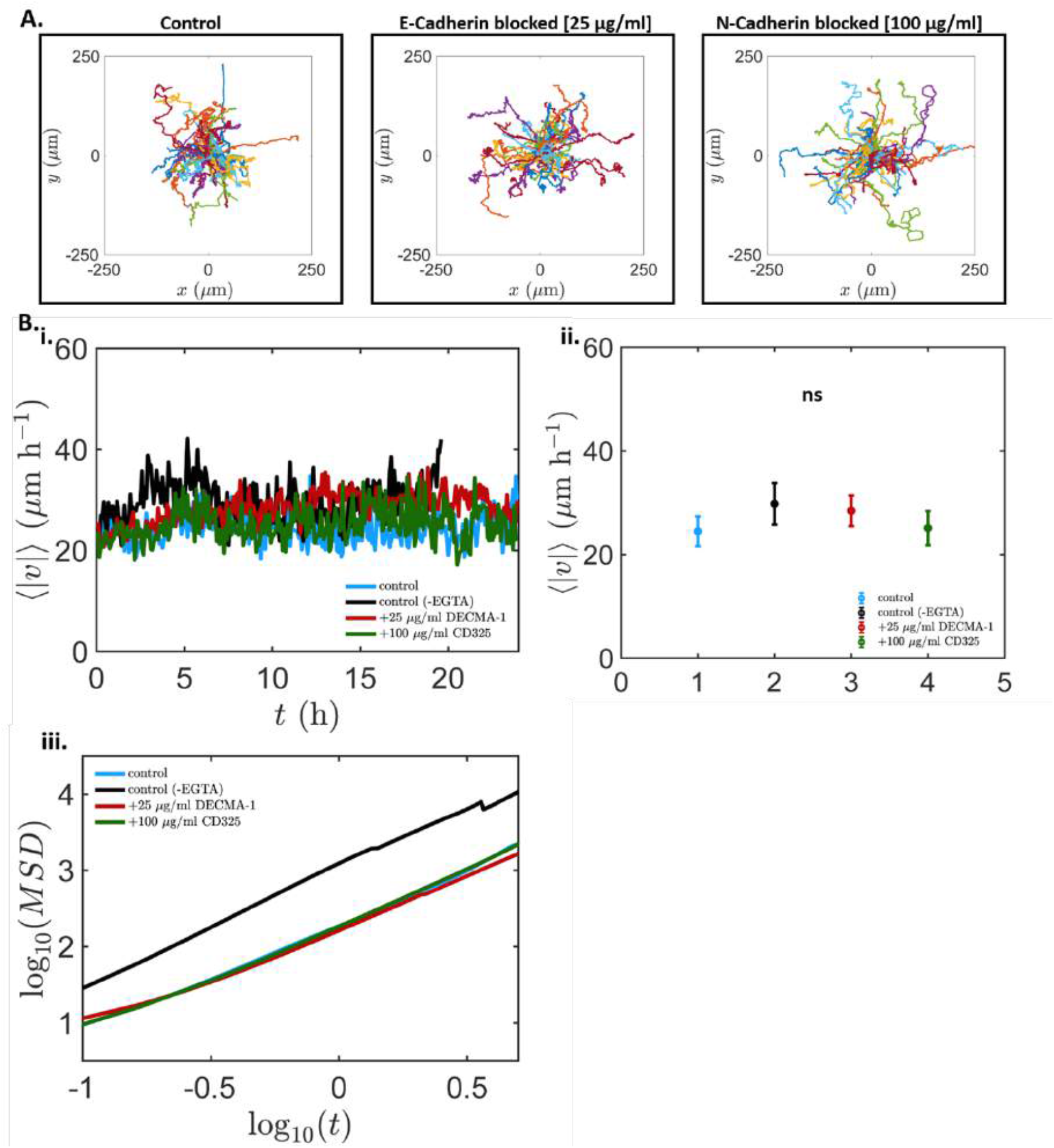
Single cell trajectories, radial velocities and SEM for single T24 cell migration in the different blocking conditions. A) Single T24 cell trajectories in the control condition and at the highest blocking antibody concentration for each cadherin type. B) i) Mean radial velocity (ur) over time and (ii) corresponding SEM graph showing no significant differences (1-way ANOVA, p>0.05) between averaged cell velocities of single cells for every condition. Average velocity of single cells was stable and was not affected by the addition of the different antibodies or EGTA pre-treatment. iii) Mean square displacement (MSD) plot showing all conditions having a 1.3 curve gradient.

**Figure S2.**
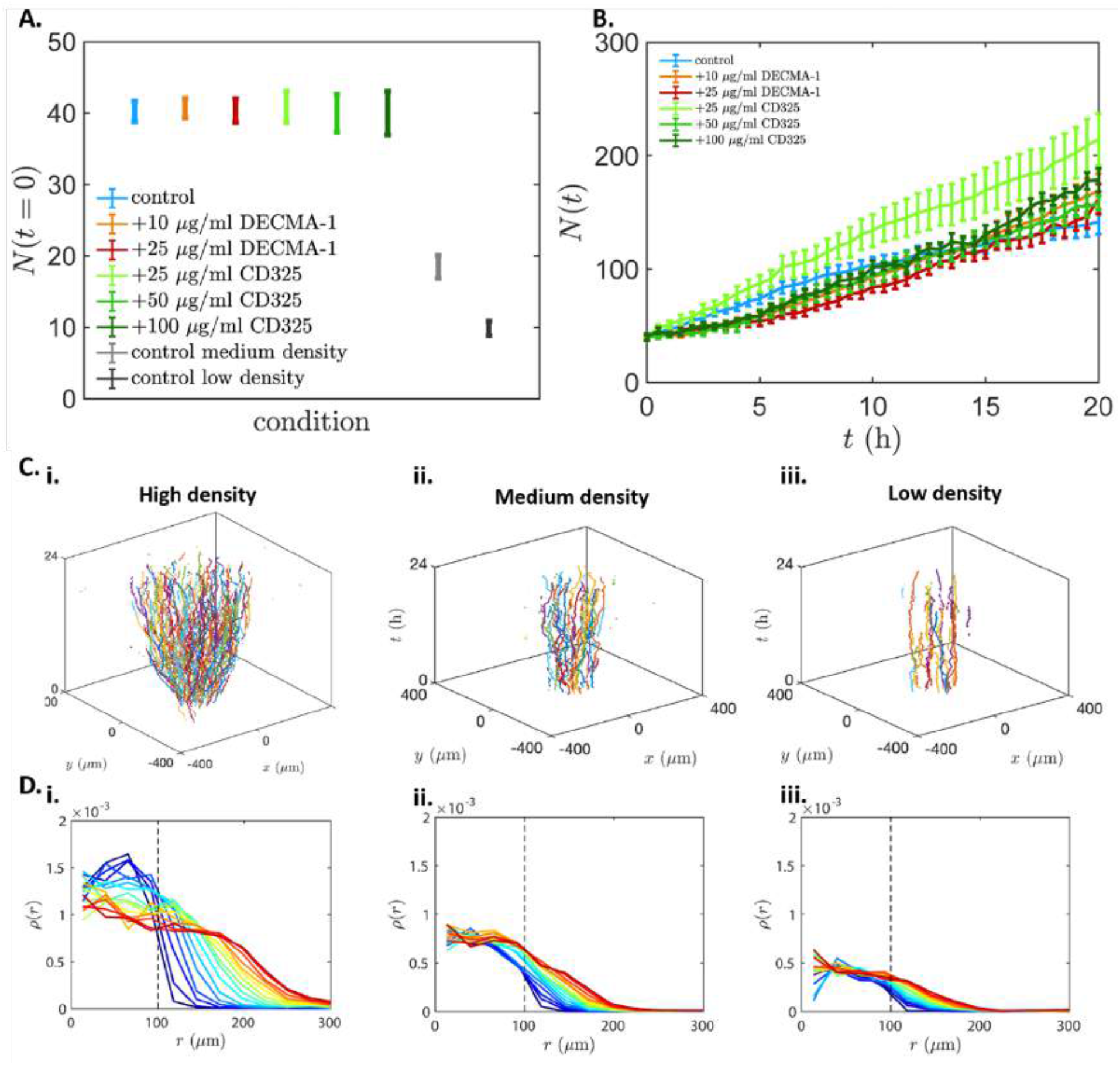
Initial number of cells of each colony for every condition and T24 cell proliferation in the different blocking conditions, followed by space diagrams and evolution of density profiles for colonies with different cell densities. A) Number of cells (t=0) for each colony per condition, color-coded according to the colony’s cell density. For every condition we ensured constant average initial cell density with an average cell number of ∼40, except medium and low cell density control conditions. B) Cell proliferation shown as the average total number of cells of all colonies for each blocking condition. In all conditions, except the 25 μg/ml N-Cadherin blocking and the combination blocking, the proliferation rate was not affected by treatment with blocking antibodies. C) Space diagram of (i) a high cell density, (ii) a medium cell density and (iii) a low cell density colony spreading up to 24h after surface activation. D) Evolution of the density profiles of (i) high cell density, (ii) medium cell density and (iii) low cell density colonies over time (blue to red) plotted as the mean of all colonies (n_high_= 12, n_medium_= 15, n_low_= 12). All curves are separated by 1.5 h intervals.

**Figure S3.**
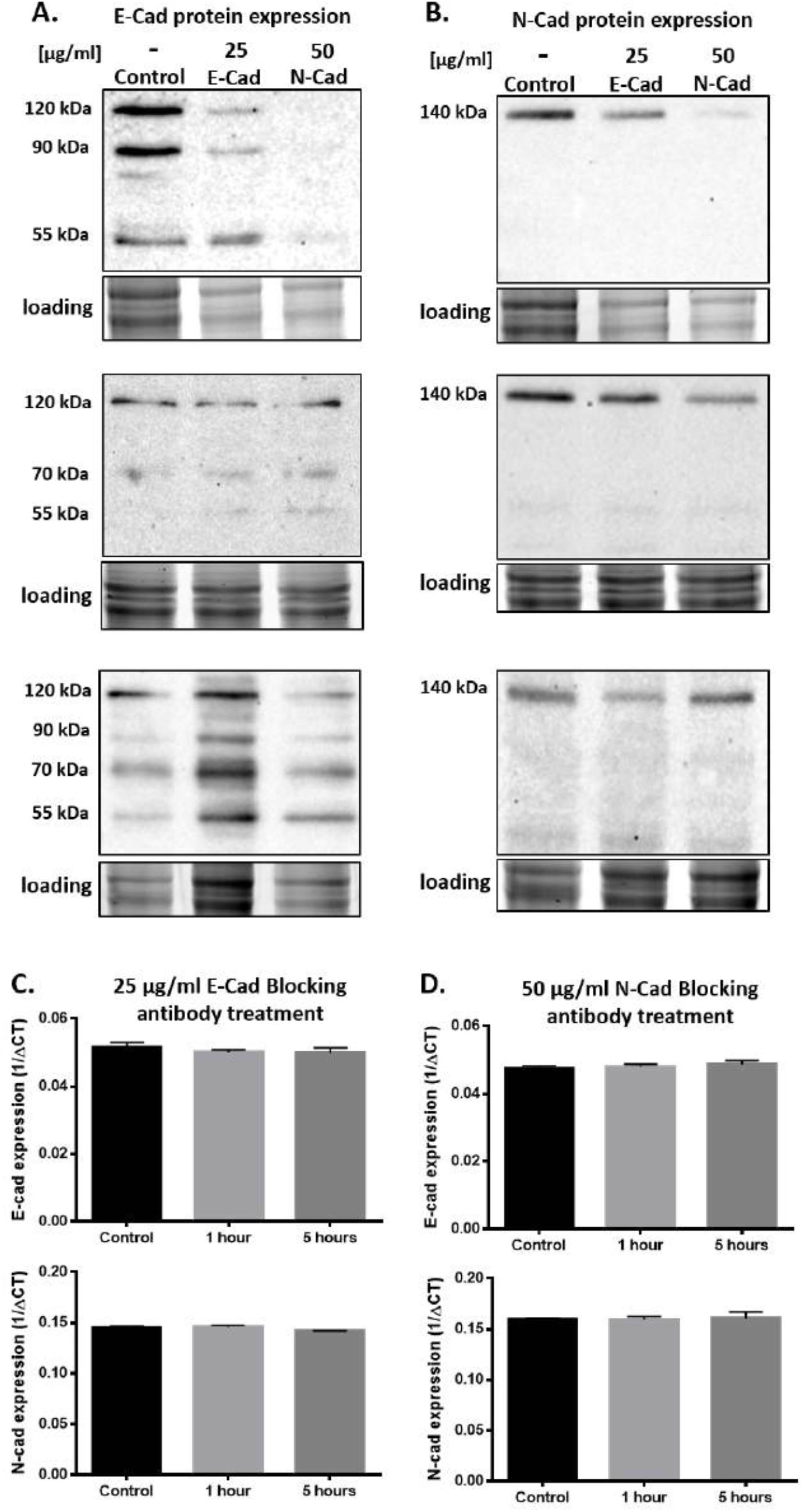
Complete Western blot triplicates and qPCR results. A) E-cadherin protein levels in untreated (control) or treated T24 cells with 25μg/ml E-Cadherin or 50μg/ml N-Cadherin blocking antibody after 24 hours. WB triplicates show the different band sizes occurring and the accompanying protein loading, with the 120kDa being the functional protein observed in all three WBs. B) N-cadherin protein levels in untreated (control) or treated T24 cells with 25μg/ml E-Cadherin or 50μg/ml N-Cadherin blocking antibody after 24 hours. WB triplicates show the 140kDa band size occurring and the accompanying protein loading. In all cases, to calculate protein expression levels all band intensities were calibrated according to control using the loading band intensities. C) No significant differences in gene expression levels of E- and N-Cadherin at 1h and 5h upon 25μg/ml E-Cadherin blocking antibody treatment as determined by qPCR (E-Cadherin: 1-way ANOVA F=0.5435, p=0.6068; N-Cadherin: 1-way ANOVA F= 3.519, p=0.0974). D) No significant differences in gene expression levels of E- and N-Cadherin at 1h and 5h upon 50μg/ml N-Cadherin blocking antibody treatment as determined by qPCR (E-Cadherin: 1-way ANOVA F= 0.4286, p=0.6699; N-Cadherin: 1-way ANOVA F= 0.04005, p=0.9610). In all cases the levels of N-Cadherin expression are ∼3-fold higher than the levels of E-Cadherin expression.

**Figure S4.**
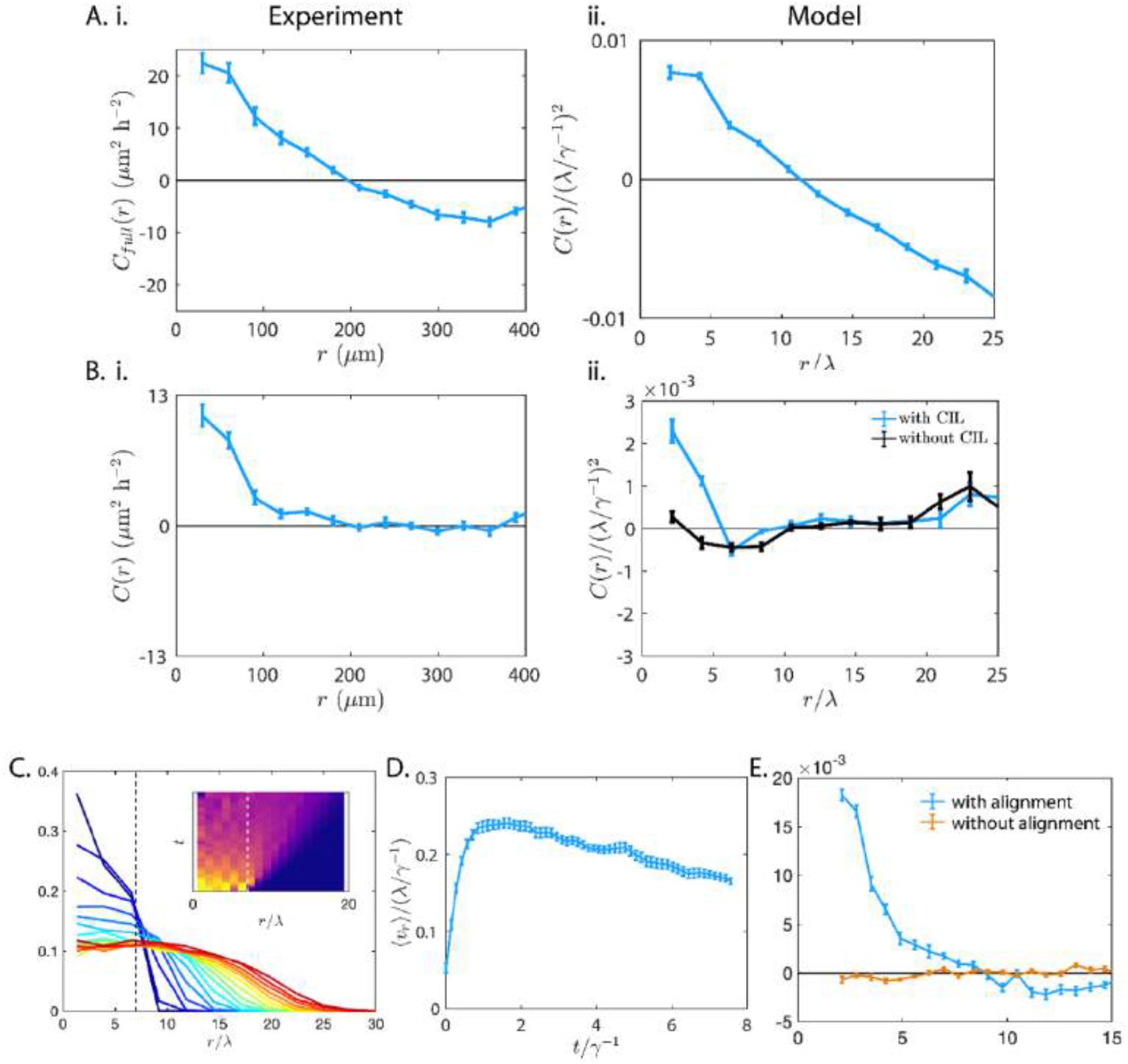
Calculation of the spatial fluctuation velocity cross-correlation function and comparison to a model with velocity alignment interactions. A) Correlation function of the full velocities for (i) experiment and (ii) model with CIL. As expected, the correlation function is initially positive, corresponding to neighboring cells on average moving in the same direction. After a distance on the order of the initial confinement radius, the function turns negative, corresponding to cells on opposite ends of the cluster moving on average in opposite directions. B) Correlation function of the velocity fluctuations, where the overall dilatation of the cluster is subtracted. For the model, corresponding curves for a simulation with CIL (blue), and without CIL interactions (black) is shown. As expected, in both cases, the negative part of the correlation due to the overall dilatation of the cluster disappears and only simulations with CIL exhibit significant fluctuation velocity correlations. C) Evolution of the density profile over time (blue to red). Inset: Kymograph of the cell density evolution. Dashed lines indicate the initial confinement radius. D) Mean radial velocity as a function of time. E) Cross-correlation of velocity fluctuations, for a model with and without velocity alignment interactions. Error bars: SEM; n=30 for all panels.

**Figure S5.**
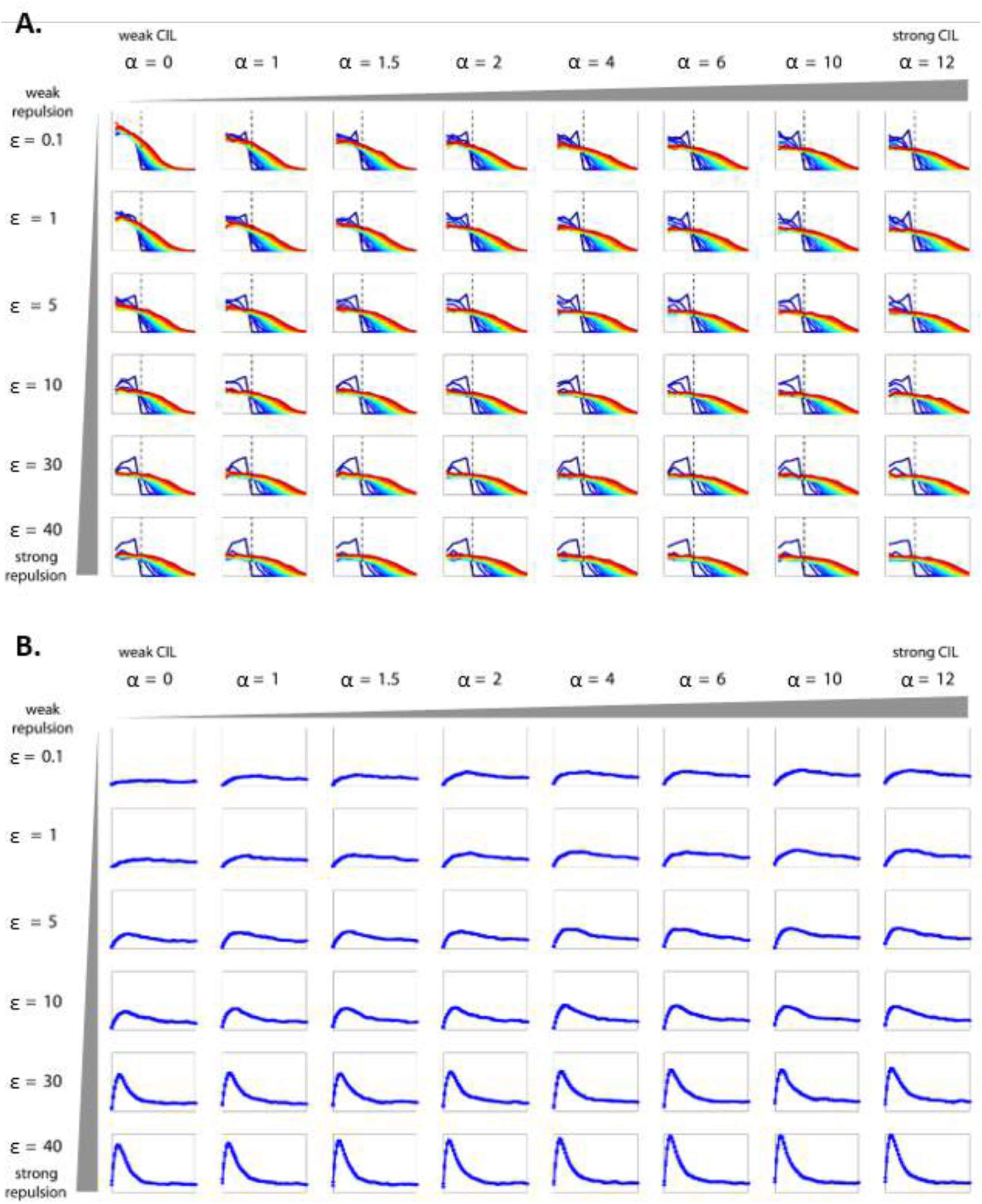
Full parameter sweep: density profiles and radial velocity curves. A) Evolution of the density profile over time (blue to red) for all parameter combinations of repulsion strength *ε* and CIL interaction amplitude *α*, averaged over n=30 clusters per condition. The profiles exhibit further spreading for larger repulsions and larger CIL amplitudes. B) Mean radial velocity as a function of time for all parameter combinations of repulsion strength *ε* and CIL interaction amplitude *α*. We generally observe larger radial velocity peaks for larger repulsions and larger CIL amplitudes. Error bars: SEM; n=30 for all panels.

**Figure S6.**
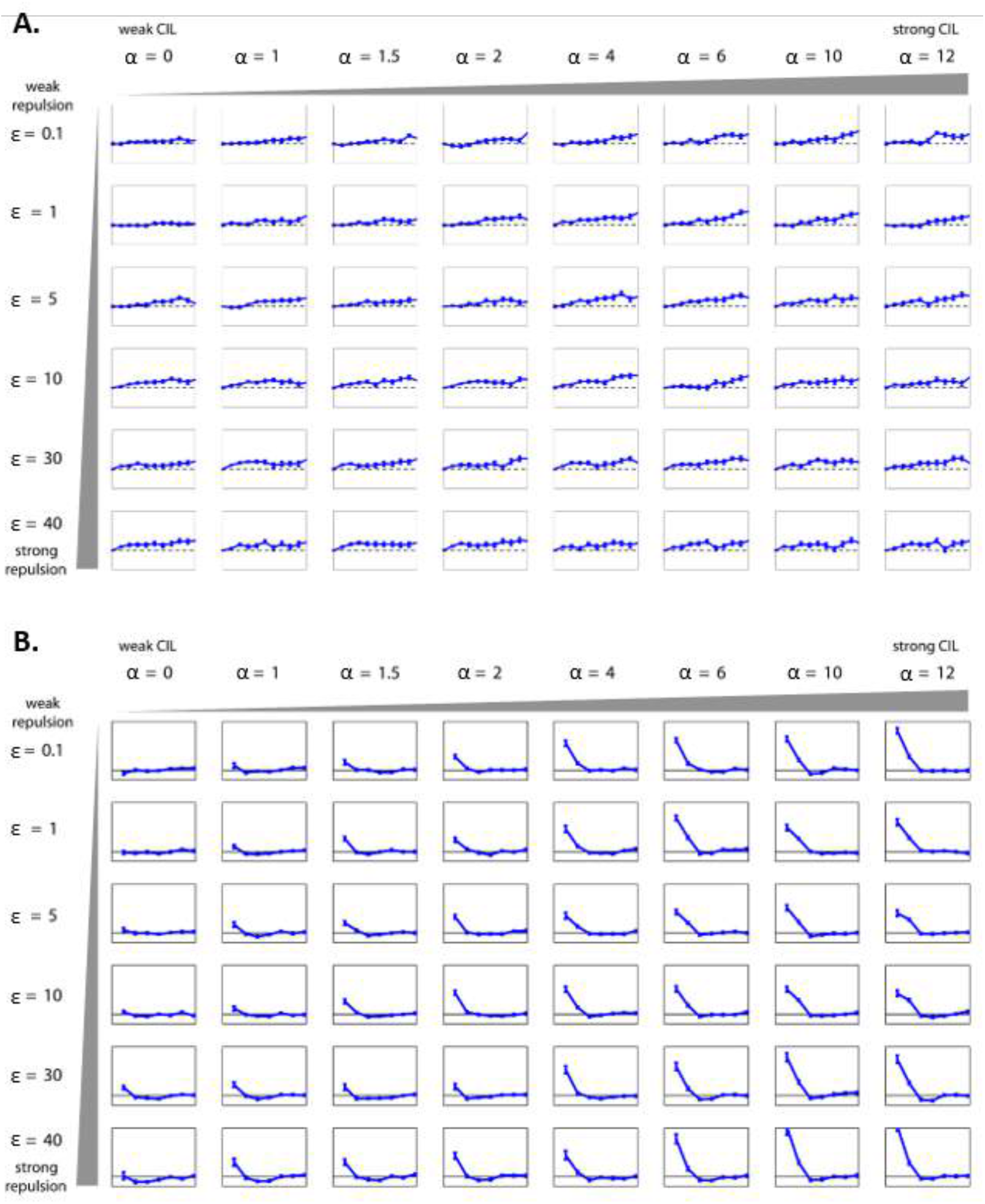
Full parameter sweep: spreading radius and velocity fluctuation cross-correlation function. A) Average distance where density has decayed to half of its value in the center of the original confinement (i.e. at r=0) for all parameter combinations of repulsion strength *ε* and CIL interaction amplitude *α*. We observe larger spreading radii for larger repulsions and larger CIL amplitudes. Error bars: SEM; n=30 for all panels. B) Cross-correlation of velocity fluctuations for all parameter combinations of repulsion strength *ε* and CIL interaction amplitude *α*. We generally observe larger radial velocity peaks for larger repulsions and larger CIL amplitudes. Error bars: SEM; n=30 for all panels.

